# Genes critical for development and differentiation of dopaminergic neurons are downregulated in Parkinson’s disease

**DOI:** 10.1101/2020.03.21.001552

**Authors:** Aditi Verma, Priya Suresh, Barathan Gnanabharathi, Etienne C. Hirsch, Vijayalakshmi Ravindranath

**Author notes:** Address for correspondence: Vijayalakshmi Ravindranath, Centre for Brain Research, Indian Institute of Science, C.V. Raman Avenue, Bangalore – 560012, India.

## Abstract

We performed transcriptome analysis using RNA sequencing on substantia nigra pars compacta (SNpc) from mice after acute and chronic 1-methyl-4-phenyl-1,2,3,6-tetrahydropyridine (MPTP) treatment and Parkinson’s disease (PD) patients. Acute and chronic exposure to MPTP resulted in decreased expression of genes involved in sodium channel regulation. However, upregulation of pro-inflammatory pathways was seen after single dose but not after chronic MPTP treatment. Dopamine biosynthesis and synaptic vesicle recycling pathways were downregulated in PD patients and after chronic MPTP treatment in mice. Genes essential for midbrain development and determination of dopaminergic phenotype such as, LMX1B, FOXA1, RSPO2, KLHL1, EBF3, PITX3, RGS4, ALDH1A1, RET, FOXA2, EN1, DLK1, GFRA1, LMX1A, NR4A2, GAP43, SNCA, PBX1, and GRB10 were downregulated in human PD and overexpression of LMX1B rescued MPP^+^ induced death in SH-SY5Y neurons. Downregulation of gene ensemble involved in development and differentiation of dopaminergic neurons indicate their critical involvement in pathogenesis and progression of human PD.

## Introduction

Parkinson’s disease (PD) is a debilitating neurodegenerative movement disorder. The cardinal clinical symptoms of Parkinson’s disease include rigidity, bradykinesia and resting tremors. PD is characterized by the loss of dopaminergic neurons in the substantia nigra pars compacta (SNpc). While it has been more than 200 years since PD was first characterized by James Parkinson^1^, lack of thorough understanding of the mechanisms that underlie disease pathogenesis has been a major obstacle in discovery of curative treatment modalities. Levodopa administration has been the mainstay for symptomatic treatment for almost sixty years^2,3^, we do not, as yet, have a disease modifying therapy that can slow down disease progression.

Familial PD is seen in less than 10% of cases, the majority are sporadic^4^. The etiology of both familial and sporadic PD is not yet clearly understood, and it is considered to be a complex, multifactorial disorder. Several risk factors have been implicated including oxidative stress, mitochondrial^5,6^ and proteasomal dysfunction^7,8^, α-synuclein aggregation^9^, neuroinflammation^10^, dopamine metabolism^11^ and iron accumulation^12,13^. However, the exact cause underlying PD pathogenesis remain poorly understood. Therefore, understanding the mechanisms that lead to neurodegeneration of dopaminergic neurons in SNpc is vital.

Transcriptomics allows us to identify dynamic changes in gene expression between different physiological states. Global gene expression analysis using microarray has been deployed to identify critical changes in gene expression in PD^14^. Many pathways, such as mitochondrial electron transport system, ubiquitin-proteasome system, apoptosis, actin cytoskeleton, inflammation and synaptic transmission have been reported to be dysregulated in PD studies^15–19^. However, microarray lacks the sensitivity to detect genes which are expressed in low levels, that is they have limited dynamic range of detection^20,21^. Moreover, there is low signal to noise ratio as well as there is an upper limit to detection of transcripts beyond which the signal gets saturated. More recently approaches using RNA sequencing (RNA-seq) offer good alternative to examine global gene expression changes in disease state.

We, therefore, used RNA seq to identify global gene expression changes in mouse model of PD following single and chronic exposure to MPTP. Further, RNA-seq was performed to study gene expression alterations in SNpc from human PD patients compared with age-matched controls.

## Results

### Gene expression changes in mouse model of PD using MPTP

RNA sequencing was performed on SNpc, dissected (Fig. 1A) from mice treated with a single dose (acute) of MPTP (30 mg/kg body weight) or daily for 14 days (chronic). While single dose of MPTP represents the first hit without substantial cell loss; about 30% loss of dopaminergic neurons is seen after 14 days of MPTP treatment^22^. Indeed, stereological examination of tyrosine hydroxylase (Th) positive neurons in SNpc revealed a 27% reduction in total number of dopaminergic neurons (Fig. 1B). Further, down-regulation of several dopaminergic genes such as, dopamine transporter (Dat), DOPA decarboxylase (Ddc) and vesicular monoamine transporter 2 (Vmat2) was observed after 14 days of MPTP treatment (Fig. 1C) but not after single exposure. Only Th was downregulated after both 1 and 14 days of MPTP treatment.

**Figure 1:**
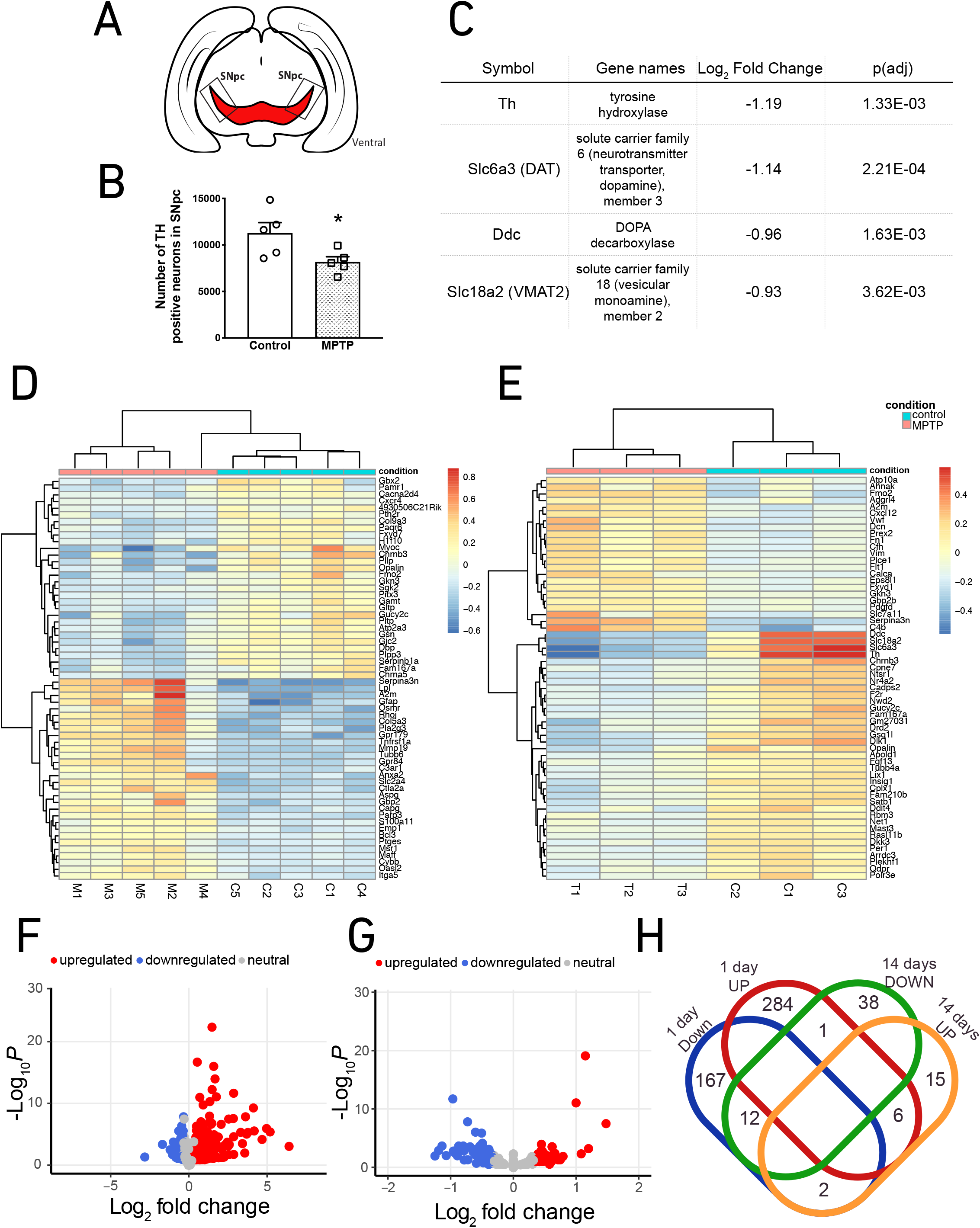
Differential gene expression in SNpc from MPTP mouse model of PD. (A) A diagrammatic representation of SNpc dissection using coronal mouse midbrain sections. (B) Stereological analysis of tyrosine hydroxylase positive neurons in SNpc after 14 days of MPTP treatment (n=5). (C) Table depicting downregulation of genes specific to dopaminergic neurons in SNpc from 14 days MPTP treated mice. (D) and (E) The top 60 significantly differentially expressed genes after 1 day and 14 days of MPTP treatment, represented through heatmaps along with their log2 fold change, respectively. (F) and (G) Volcano plots for differentially expressed genes in SNpc from 1 day and 14 days MPTP treated mice SNpc, respectively. (H) Venn diagram showing the overlap of the upregulated and downregulated genes between 1 day and 14 days MPTP treatment. Each dot represents an individual on the graphs. Comparisons between the groups were done using unpaired Student’s t test. Data are presented as mean ± SEM. * indicates p<0.05.

Differential gene expression analysis of RNA-seq data from mouse SNpc after 24 hours following single exposure to MPTP revealed significantly (adjusted p <0.05) altered expression of 472 genes, 291 of which were up-regulated and 181 were down-regulated (Fig. 1D and 1F). However, analysis of data from mouse SNpc after 14 days of MPTP treatment showed significant (adjusted p <0.05) altered expression of 74 genes, 23 of which were up-regulated and 51 were down-regulated (Fig. 1E and 1G). Thus, fewer genes were perturbed as the disease progressed and cell loss continued. As evident from the Venn diagram (Fig. 1H), there was little overlap between the two sets suggesting the involvement of different molecular pathways during early and later stages of MPTP treatment. Further, there was greater perturbation in terms of number of differentially expressed genes 24 hours after single dose of MPTP, which could be due to the adaptive response to the toxin. A complete list of the differentially expressed genes is provided in the Supplementary Tables S1 and S2.

### Perturbation of different pathways seen at early and late stage of MPTP mouse model of PD

Identification of gene ontology (GO) pathways including biological processes (BP), molecular function (MF), and cellular components (CC), which were populated by genes that were significantly up or downregulated in SNpc after single exposure to MPTP revealed an upregulation of pathways, such as innate immune response, inflammatory response, response to cytokine, cellular response to tumor necrosis factor, apoptotic processes, extracellular exosome, MAPK cascade, regulation of cell proliferation, aging and dendrite and a downregulation of pathways, such as nervous system development and sodium channel regulator activity (Fig. 2A and Supplementary Table S3). A similar analysis for the RNA-seq data from 14 days MPTP mouse SNpc did not show perturbation of innate immune response or inflammatory response. However, upregulation of pathways involved in functioning of plasma membrane, extracellular space, extracellular exosome and downregulation of pathways involved in functioning of axon, response to nicotine, response to toxic substances, response to hypoxia, neuron projection, cytoplasmic vesicle, regulation of dopamine metabolic process, regulation of sodium ion transport, synapse, neuronal cell body and dendrite (Fig. 2B and Supplementary Table S4).

**Figure 2:**
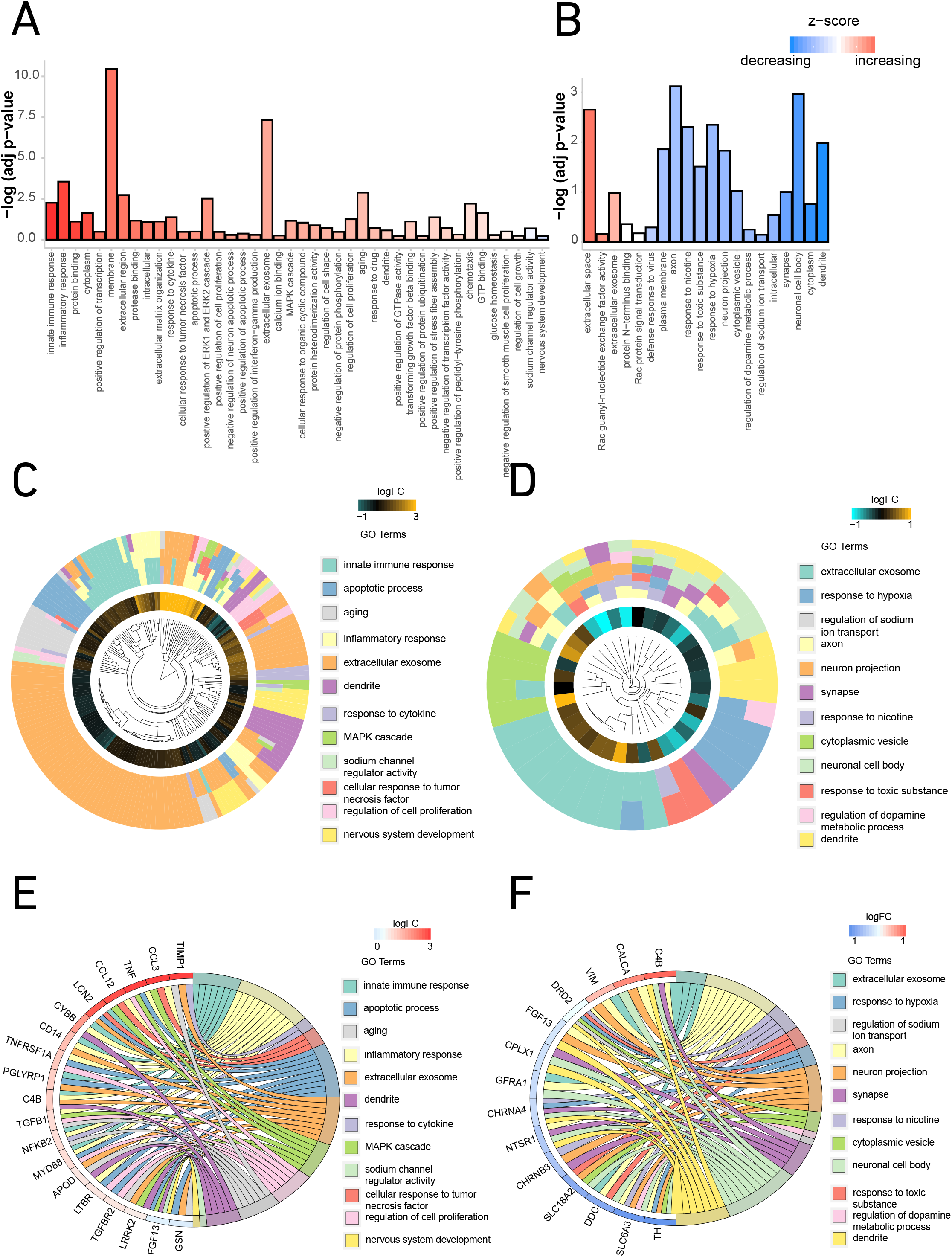
Differentially expressed pathways enriched in SNpc from MPTP mouse model of PD. (A) and (B) Bar diagrams for functional clusters derived using the Database for Annotation, Visualization and Integrated Discovery (DAVID) for significantly differentially expressed genes in 1 day and 14 days MPTP mouse model of PD, respectively. (C) and (D) Significantly, differentially expressed genes and their distribution into pathways hierarchically clustered according to functional categories along with the adjusted p values for 1 day and 14 days MPTP treated mice SNpc using circular plots, respectively. The innermost space represents the hierarchical clustering of the genes while the inner circle represents the log_2_ fold change. (E) and (F) Chord diagrams for pathways depicting the significant genes that belong to at least three different pathways with their log_2_ fold change for 1 day and 14 days MPTP treatment in mice, respectively.

The circular plots for acute (Fig. 2C) and chronic (Fig. 2D) MPTP treatment shows hierarchical clustering of the significant genes based on functional categories and how they populate these categories along with their log_2_ fold change. Further, chord diagrams for differentially expressed pathways and their resident genes after single dose (Fig. 2E) and 14 days (Fig. 2F) of MPTP treatment show the significantly differentially expressed genes that populate at least three different pathways and their log2 fold change.

The early stage of MPTP mouse model thus presents a canvas of molecular pathways that is quite dissimilar to what is observed at late stage of the model. It is also intriguing to note the absence of differentially expressed genes related to innate immunity and inflammatory response in SNpc from mouse treated with MPTP for 14 days, wherein about 30% of dopaminergic neurons are lost.

### Gene expression changes in SNpc from human PD

The anatomical landmarks used for dissection of SNpc are depicted in Fig. 3A. The top 30 differentially expressed genes are tabulated in Fig. 3B. Differential gene expression analysis through DESeq2 revealed significantly (adjusted p <0.05) altered expression of 367 genes, of which 103 were up-regulated and 264 were down-regulated in SNpc of human PD brain (Fig. 3C and 3D). Complete list of differentially expressed genes is provided in Supplementary Table S5. Markers of dopaminergic neuron phenotype such as DAT, VMAT2, DDC, DRD2 and TH were significantly downregulated (Fig. 3E), which served as a positive indication of degeneration of dopaminergic neurons in SNpc in these samples. Components of the ubiquitin-proteasome pathway such as, PSMD12, UBE2T, UBFD1 and UCHL1 were down-regulated in the human PD. Of the genes that were down-regulated, UCHL1 (also known as PARK5) has been associated with familial PD^23^. Dysregulation of the ubiquitin-proteasome pathway is one of the well-studied mechanisms implicated in the pathogenesis of PD^24^.

**Figure 3:**
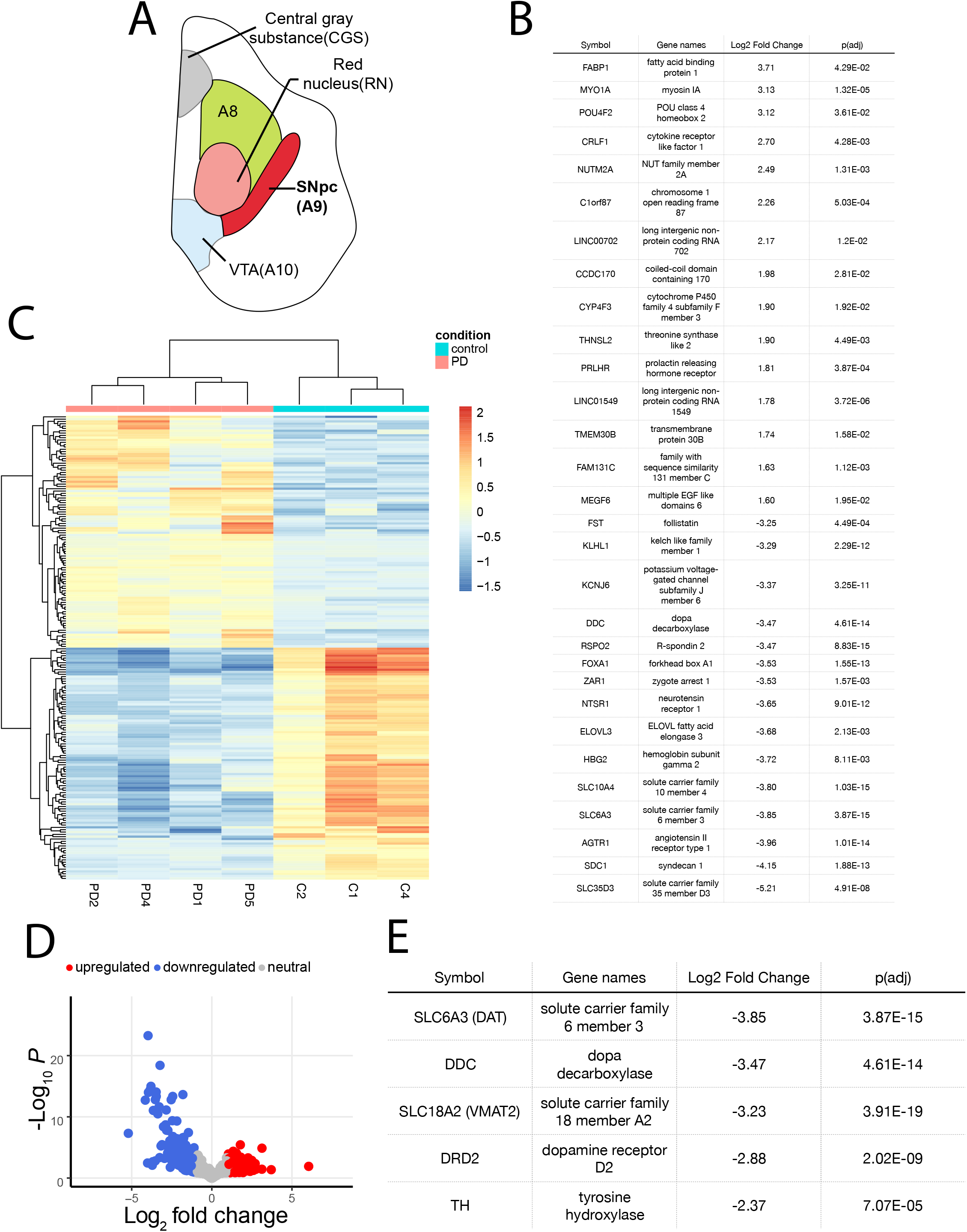
Differential gene expression in SNpc from human autopsy tissue samples. (A) Graphical representation of human midbrain nuclei, namely central gray substance (CGS), red nucleus (RN), peri and retro rubral dopaminergic cell group (A8), substantia nigra pars compacta (SNpc – A9)), ventral tegmental area (VTA – A10). (B) Differential gene expression in PD-from human autopsy tissue. The list displays the top 30-upregulated and downregulated genes in SNpc from PD autopsy tissue. The log_2_ fold change and adjusted p value are also provided along with gene names. (C) Heatmap depicting differentially expressed genes in SNpc from human PD (D) Volcano plot representing the relative numbers of upregulated and downregulated genes. (E) Downregulation of dopaminergic neuron markers in SNpc from autopsy tissue from PD patients.

Neuron specific genes such as MAP2, RBFOX3 and TUBB3 were unchanged in human PD. Further, the mRNA expression of β-actin and glyceraldehyde 3-phosphate dehydrogenase (GAPDH), as assessed using qRT-PCR, remained unchanged as seen after normalization with ribosomal protein L13 (RPL13) in human PD (Supplementary Fig S1)

### Pathway analysis reveals down regulation of dopaminergic neuron differentiation factors in human PD

Pathway analysis, performed using the significantly differentially expressed genes, showed downregulation of functional clusters, such as those involved in oxidation-reduction process, neuron projection, synaptic vesicle recycling, regulation of synaptic plasticity, dopamine biosynthetic process, dopaminergic neuron differentiation and locomotory behavior (Fig. 4A and Supplementary Table S6). We also clustered significantly expressed genes according to functional categories (Fig. 4B). Further, the chord diagram shows distribution of differentially expressed genes in pathways with their adjusted p values (Fig. 4C). An interesting set of genes that were perturbed significantly as highlighted is of those that belong to dopaminergic neuron differentiation as also those that define midbrain dopaminergic neuron development and maintenance. These include FOXA1, RSPO2, KLHL1, LMX1B, EBF3, PITX3, RGS4, ALDH1A1, RET, FOXA2, EN1, DLK1, GFRA1, LMX1A, NR4A2, GAP43, SNCA, PBX1 and GRB10 (Fig. 4D).

**Figure 4:**
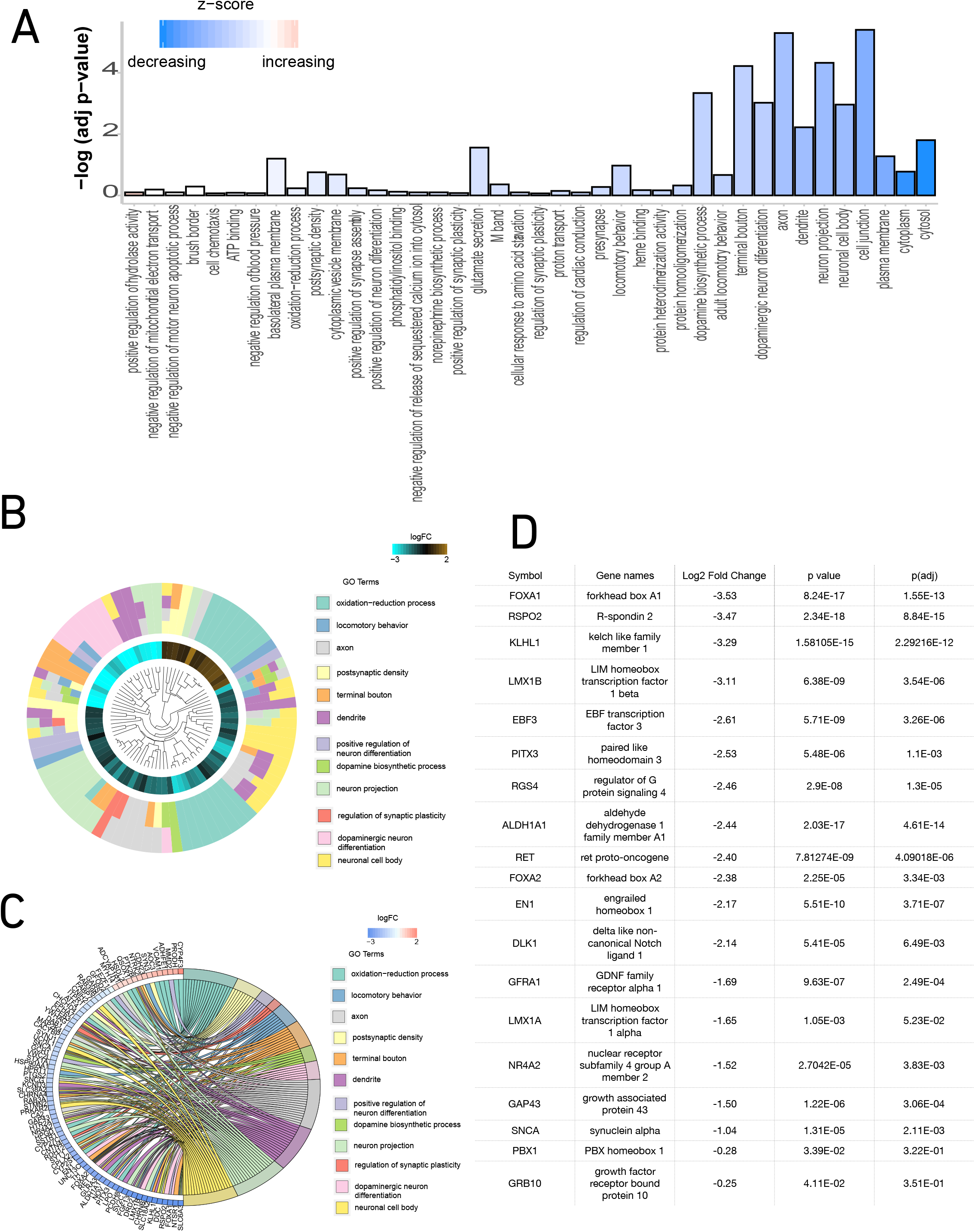
Differentially expressed pathways enriched in SNpc from human autopsy tissue samples. (A) Bar diagram of the differentially regulated functional clusters as derived using DAVID from the significantly differentially expressed genes. (B) Circular plot for the significant genes enriched in pathways where the innermost space represents the hierarchical clustering of the genes while the inner circle represents the log_2_ fold change. (C) Significantly differentially expressed genes that belong to at least two functional clusters and their log2 fold change as represented using a chord diagram. (D) Table of genes that are involved in development, differentiation and maintenance of midbrain dopaminergic neurons with their log_2_ fold change, p value and adjusted p value.

We validated the downregulation of the genes belonging to the pathway related to differentiation of midbrain dopaminergic neurons, viz. GAP43, RET, LMX1B and EN1 in SNpc from PD autopsy tissue using qRT-PCR. Evaluation of TH mRNA was not performed since four different isoforms exist in the human brain^25,26^ while a single TH isoform is present in the mouse brain^27^, creating difficulty in choosing a single TH sequence as a suitable reference marker for qRT-PCR in human tissue. Downregulation of DAT, a marker for dopaminergic neurons was seen (Fig. 5A), which is expected following dopaminergic neuronal death in SNpc. This is indeed an interesting observation considering that genes involved in differentiation of dopaminergic phenotype are indeed important for its maintenance and their downregulation could impact the health of dopaminergic neuron.

**Figure 5:**
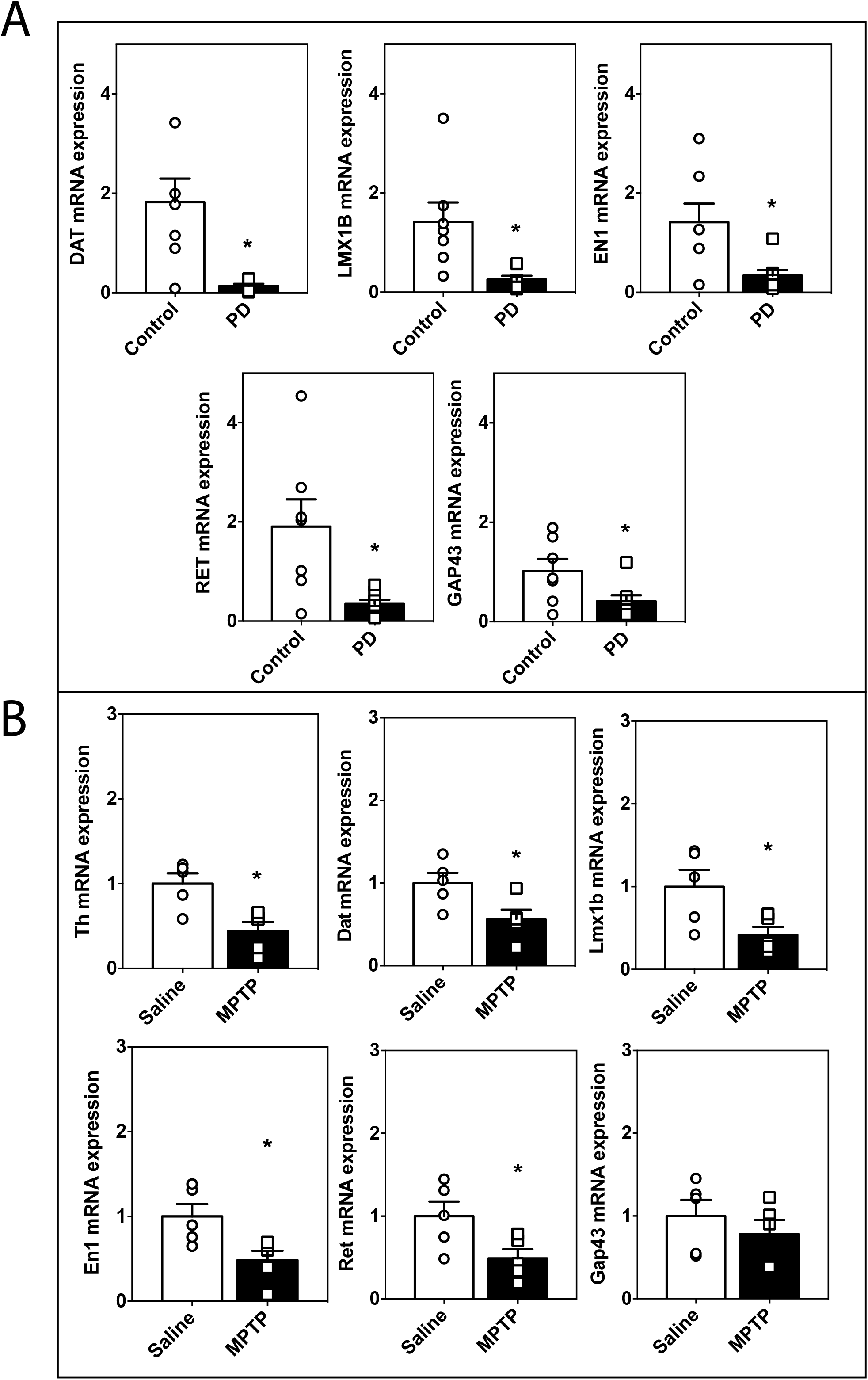
Validation of down-regulation of genes involved in development and differentiation of midbrain DA neurons through qRT-PCR. (A) Relative level of gene expression of DAT, LMX1B, EN1, RET and GAP43 in on human autopsy SNpc as compared to age-matched controls as quantified using qRT-PCR. (B) Significant downregulation of Th, Dat, Lmx1b, En1 and Ret but not of Gap43 observed in SNpc from C57BL/6 mice treated with MPTP (s.c.) for 14 days. Each dot represents an individual on the graphs. Comparisons between the groups were done using unpaired Student’s *t* test. n=5-8; data has been presented as mean ± SEM. * indicates p<0.05.

We further validated the down-regulation of genes using an alternate method of normalization, that is, Cufflinks and also after regressing the data for age, post mortem interval (PMI) and RIN using DESeq2 (Supplementary Tables S7 and S8) and found that majority of the selected genes were significantly downregulated through both the analyses. Though the downregulation of GAP43 does not have a significant adjusted p value but only a significant p value, it was significantly down-regulated in the qRT-PCR experiments.

The RNA-seq data from mouse SNpc following MPTP treatment for 14 days did not show significant adjusted p values for Lmx1b, En1 and Ret even though the raw reads had decreased by 39, 18 and 37%, respectively. However, these genes were significantly downregulated when examined using qRT-PCR (Fig. 5B). The downregulation of these genes in mouse model of PD demonstrate that the dysregulation of this pathway occurs both in mouse and human.

Protein-protein interactions that can potentially result from co-expression changes in the genes involved dopaminergic neurons differentiation, is depicted in Fig. 6A using String database^28^. Further, mapping these interactions based on existing data (Fig. 6B) revealed LMX1B as an important gene upstream to the expression of many genes involved in dopaminergic midbrain neuron development, differentiation and maintenance. Thus, we see that the whole ensemble of genes (LMX1B, FOXA1, RSPO2, KLHL1, LMX1B, EBF3, PITX3, RGS4, ALDH1A1, RET, FOXA2, EN1, DLK1, GFRA1, LMX1A, NR4A2, GAP43, SNCA, PBX1 and GRB10) that are essential for dopaminergic neuron development, differentiation and maintenance are downregulated in human PD, including those that are downstream of LMX1B. Therefore, we overexpressed LMX1B and assessed its effect on MPP^+^ mediated cell death.

**Figure 6:**
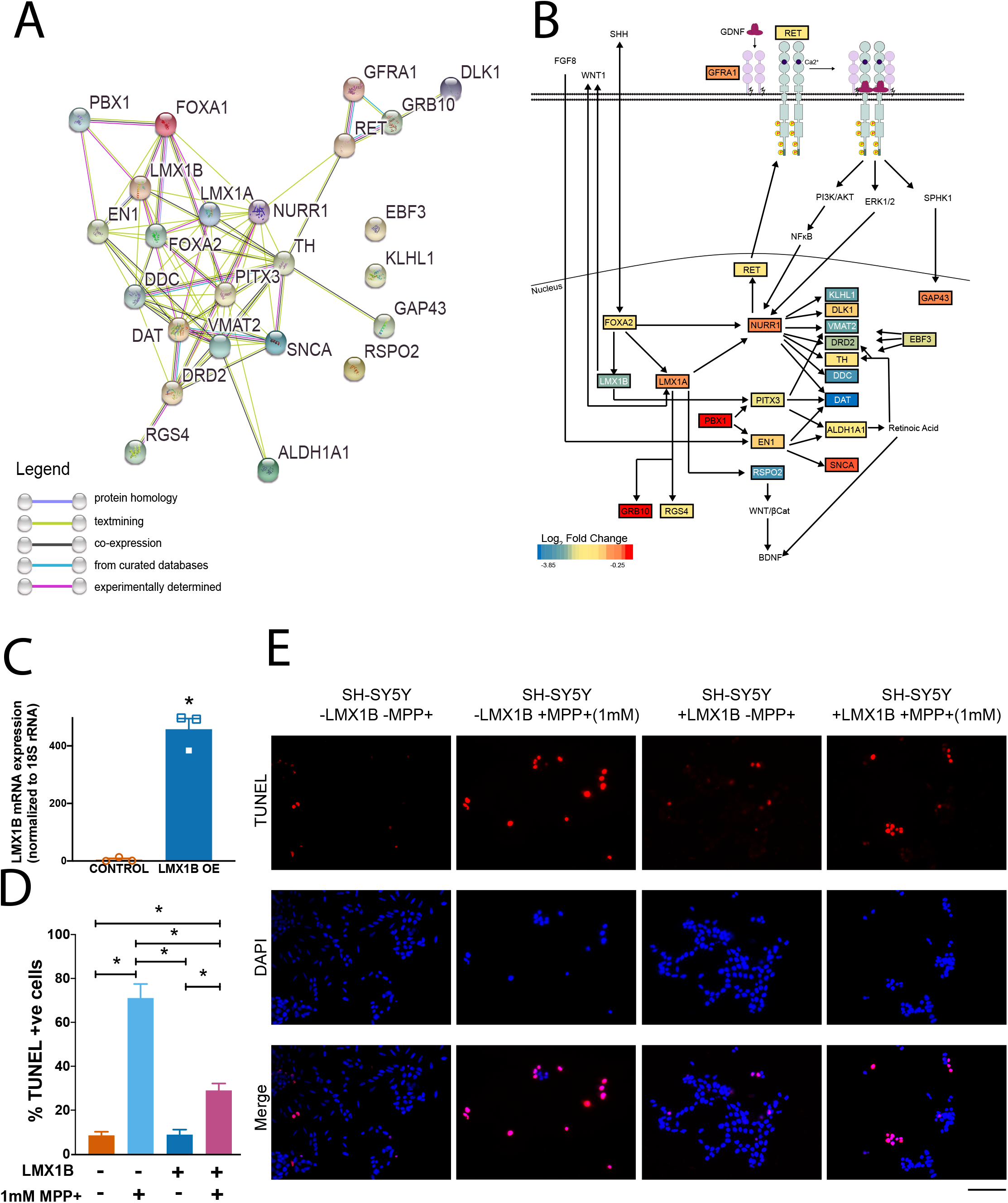
LMX1B overexpression rescues SH-SY5Y from MPP^+^ toxicity. (A) String analysis of significantly perturbed midbrain dopaminergic neuron development, differentiation and maintenance genes. (B) Summary of role of genes involved in development and maintenance of SNpc dopaminergic neurons that were differentially expressed in human PD as derived from literature. (C) Confirmation of increased gene expression of LMX1B in SH-SY5Y cells after transfection with LMX1B overexpression (LMX1B OE) plasmid as assessed using qRT-PCR. (D) Quantification of percentage of TUNEL positive cells as normalized to the total number of DAPI positive cells. n=28-30 images, 8-10 images taken per coverslip taken over three independent coverslips. (E) Representative images of SH-SY5Y neurons depicting the presence of TUNEL positive cells (Red) and DAPI positive cells (Blue) after overexpression of LMX1B and treatment with 1mM MPP^+^. Comparison between the groups was done using unpaired Student’s *t* test. Data represented as mean ± SEM. One-way ANOVA with Tukey’s test for multiple comparison was performed to compare the different groups. * represents p<0.05. Scale bar represents 100μm.

### Lmx1B overexpression rescues SHSY-5Y from MPP^+^ toxicity

Recombinant human LMX1B was overexpressed in SH-SY5Y neuronal cell line, as confirmed using qRT-PCR (Fig. 6C), which were then exposed to parkinsonism-inducing toxin, MPP^+^. We observed that while exposure to MPP^+^ led to significant death in SH-SY5Y cells as measured using TUNEL assay, overexpression of LMX1B led to significant reduction in the percentage of TUNEL positive cells (Fig. 6D and 6E). Our results, therefore, show that LMX1B is capable of rescuing SH-SY5Y cells from MPP^+^ mediated cell death.

## Discussion

In this study, global changes in the transcriptome were studied in precisely dissected substantia nigra pars compacta (SNpc) from MPTP mouse model of PD after acute and chronic MPTP treatment using RNA sequencing. Further, RNA sequencing was also performed on SNpc from PD patients and age-matched controls. A single dose of MPTP resulted in a larger number of differentially expressed genes in mouse SNpc as compared to 14 days treatment with MPTP. Gene expression changes in SNpc from mice after single exposure to MPTP resulted in dysregulation of pathways related to immune and inflammatory system. However, after 14 daily doses of MPTP and in human PD, these pathways were not enriched. While Riley and colleagues have indeed reported dysregulation of several inflammatory pathways^29^, it is essential to note that the RNA-seq in their study was performed on substantia nigra which would include contribution from the GABA-ergic neurons of pars reticulata. The present study, on the other hand, evaluates gene expression changes in specifically substantia nigra pars compacta, which could explain the observed difference. Further, a recent transcriptomic study describes gene expression alteration in SN from the brain of PD patients in relation to Braak staging^30^. Using co-expression analysis, Keo and colleagues have reported the dysregulation of genes related to dopamine synthesis, microglia, immune system, blood-oxygen transport, and endothelial cells. However, in the present study, when we analyzed only the significantly differentially expressed genes, we were able to identify the downregulation of an ensemble of genes involved in development and differentiation of midbrain dopaminergic neurons as a novel pathway involved in PD pathogenesis. We, further, confirmed downregulation of these genes using qRT-PCR and validated the protective effect of overexpression of an upstream regulator of this pathway namely, LMX1B, against MPP^+^ induced cell death in SHSY-5Y cells.

When we compared the transcriptomic changes observed in 14 days MPTP mouse SNpc with human PD, we found similarity in terms of downregulation of the common genes involved in dopaminergic phenotype maintenance. While few dysregulated pathways are common between MPTP model and human PD, the differences are predominant reinforcing the limited overlap of the molecular pathways involved in neurodegeneration.

Two striking observations of this study are the lack of enrichment of genes involved in inflammatory and immune response and the downregulation of an entire ensemble of genes in SNpc from PD subjects that are well characterized for their role in maintenance of dopaminergic phenotype, viz. FOXA1, RSPO2, KLHL1, LMX1B, EBF3, PITX3, RGS4, ALDH1A1, RET, FOXA2, EN1, DLK1, GFRA1, LMX1A, NURR1, SNCA, PBX1 and GRB10 in human PD^31–38^. The downregulation of these genes could presumably affect the expression of genes that define the dopaminergic phenotype, such as DAT, TH, DRD2 and DDC. GAP43 has been implicated in neurogenesis and its role in survival or maintenance of dopaminergic neurons is not clear^39,40^.

A summary of the functions of genes involved in development, differentiation and maintenance of dopaminergic neurons is presented in Fig. 6B. Sonic hedgehog (SHH), WNT1 and FGF8 are important early genes that induce the expression of FOXA2, LMX1A, LMX1B and EN1 during the earliest stages of midbrain dopaminergic neuron development, which further drive the expression of NURR1 and PITX3 and thereby lead to expression of TH, VMAT2, DRD2, DDC, DAT and ALDH1A1 that define the dopaminergic neuron phenotype^41–47^. Further, studies have implicated the role of genes such as RGS4, GRB10 and RSPO2 that are downstream LMX1A in midbrain dopaminergic neurogenesis and differentiation^48,49^. Similarly, there is evidence that EBF3 may lie upstream of NURR1 and PITX3^50^ and PBX1 is upstream of PITX3 and EN1^51–53^ and may play an important role in dopaminergic neuron development. Moreover, the expression of NURR1 and PITX3 is induced by GDNF/RET signaling and has been shown to be important for protection of dopaminergic neuron^37,54^. It is also important to note that several studies have emphasized on the strong co-ordination among the members of this pathway such as NURR1 and FOXA2, PITX3 and EN1, NURR1 and EN1 and FOXA2 and LMX1A and B^51,52,55–57^.

While these genes are indispensable for the development of midbrain dopaminergic neurons, fewer studies have assessed their function in the adult SNpc. EN genes have been reported to be essential for survival of midbrain dopaminergic neurons and regulate the expression of α-synuclein (SNCA)^58^. LMX1A/B have been shown to be important for maintenance of dopaminergic neuron phenotype through their roles in regulating normal autophagic-lysosomal pathway^59^ and mitochondrial functions^60^.

In the present study, we have observed the concomitant down-regulation of majority of the genes important for development and maintenance of dopaminergic neurons, which not only emphasize the role that this pathway plays in pathogenesis and progression of PD but also the interdependence of these genes in dopaminergic neuron survival. However, how these genes are downregulated so specifically in SNpc from PD subjects and their roles in the adult SNpc needs to be addressed. Importantly, we demonstrated that the overexpression of LMX1B, which is an upstream regulator of the midbrain dopaminergic neuron differentiation pathway, is able to rescue cell death as a result of MPP^+^ toxicity in the SH-SY5Y neuron cultures suggesting that a revival of this pathway could potentially rescue neurodegeneration in PD.

While greater emphasis has been given to addressing the motor dysfunctions observed in PD, management of non-motor symptoms of PD including sleep and autonomic dysfunction, and psychiatric symptoms is increasingly becoming a larger burden. Interestingly, review of genome wide association studies (GWAS) revealed the association of ALDH1A1^61^ and DRD2^62^ with schizophrenia. DRD2 is also associated with alcohol consumption^63^ and sleep deprivation^64^. NURR1^65^ and LMX1A^66^ are associated with alcohol consumption while DAT^67^ and PBX1^68^ are associated with sleep duration. Further RET has been associated with cannabis dependence^69^ and smoking behavior^70^. Thus, we see that several of the genes dysregulated in PD are associated with addictive behavior and impact a variety of behavioral dysfunctions which co-occur with PD. The impact of downregulation of the genes as we see in the present study would potentially have far-reaching effects beyond motor symptoms.

In conclusion, we demonstrate for the first time that an ensemble of 19 genes that govern and regulate the development of midbrain and emergence of dopaminergic phenotype in SNpc neurons are markedly downregulated in Parkinson’s disease. These not only include genes that are downstream of LIM homeobox transcription factor, LMX1B (PITX3 and ALDH1A1) and LMX1A (GRB10, RGS4, RSPO2, NURR1, KLHL1, DLK1 and RET), but also other genes that are upstream such as, FOXA2 and RET and genes regulated by FGF8 such as EN1; indicating that these genes are potentially important for the maintenance of dopaminergic neurons. Interventions that could potentially upregulate the key genes that lie upstream in this cascade could protect dopaminergic neurons and thus alter the progression of the disease. Thus, identification of this ensemble offers a new window for discovery of disease modifying therapies for PD.

## Materials and Methods

### Animals and MPTP dosing

C57BL/6J male mice (3-4 months; 25-30 grams), procured from Central animal facility of Indian Institute of Science were used for all animal experiments. Animals were housed in groups of five and had access to pelleted diet and water, *ad libitum*. For 1-methyl-4-phenyl-1,2,3,6-tetrahydropyridine hydrochloride (MPTP) mouse model of dopaminergic loss^71^, MPTP (Sigma-Aldrich Cat# M0896) dissolved in normal saline was given at a dose of 30 mg/kg body weight subcutaneously to mice as a single injection or daily for 14 days. MPTP injections were carried out in an isolated clean air room in the Central Animal Facility, Indian Institute of Science, Bangalore. Adequate safety precautions were taken for proper handling of MPTP during preparation and injection and for disposal of materials and samples contaminated with MPTP. The controls were injected with an equivalent volume of normal saline. Animals were sacrificed 24 hours after the last MPTP dose.

### Preparation of brain coronal sections for histochemical analysis

After 14 days of MPTP treatment, animals were anesthetized in an enclosed chamber and were then perfused transcardially with 4% (w/v) paraformaldehyde (PFA) that served as a fixative. The mouse brains were then carefully removed and post-fixed in 4% (w/v) PFA for 12 h at 4°C, followed by transfer to 30% (w/v) sucrose in 1X PBS at 4°C for ~36-48 h. The brains were then embedded in tissue freezing medium (Leica Microsystems Nussloch GmbH Cat# 0201 08926). Serial coronal sections (25 μm thickness) were cut through the midbrain region of the brain.

### TH immunohistochemistry

Serial coronal sections preserved in Tris buffered saline (TBS) with sodium azide (0.005% w/v) were used for IHC. Sections were washed twice with TBS (5 min) and incubated with blocking solution (3% (w/v) bovine serum albumin (BSA), 0.1% (v/v) Triton-X 100, and 3% (v/v) normal horse serum in TBS) for 1 hour at room temperature followed by incubation in rabbit anti-TH (Millipore Cat# AB152, RRID: AB_390204) (1:500) in TBS-Tween 20 (TBST) for 12 h at 4°C. Further, sections were given four washes with TBST and incubated in donkey anti-rabbit IgG H+L (Alexa Fluor^®^ 594) (ThermoScientific A-21207, RRID: AB_2556547) (1:1000) in TBST for 1 hour. The sections were then washed and mounted in VECTASHIELD^®^ mounting medium with DAPI (Vector Laboratories Cat# H-1200) on glass slides.

### Stereological analysis on histological data

Analysis of total number of TH positive neurons in SNpc was performed for every sixth coronal section through the midbrain using StereoInvestigator^®^ by MBF Bioscience. All data were quantified in an experimentally blind manner.

The sections were imaged on an Olympus microscope using the StereoInvestigator^®^ (MBF Bioscience) software to obtain tile scans of unilateral SNpc across z (vertical axis) at a magnification of 40X. Counting was performed in an experimentally blind manner, offline using the optical fractionator probe^72,73^ using 60 × 60 μm counting frame at *x* =150 μm, *y* = 150 μm intervals from a random start point. A guard zone of 2.5 μm, and a probe depth of 20 μm were used. The coefficient of error was below 0.1 in all animals studied.

### Mouse brain SNpc dissection

MPTP treated mice were decapitated following cervical dislocation post 24 hours of the last dose of MPTP in the treatment paradigm following cervical dislocation. Mouse brain was placed on mouse brain matrix (Ted Pella, Inc Cat# 15050) and 1mm thick slices of the brain were obtained for the dissection of SNpc, under cold and sterile conditions. Then, SNpc was dissected out from these slices under stereomicroscope using anatomical markers.

### Human brain tissue dissection

For the dissection of SNpc, slides containing cryostat-cut sections of flash frozen human autopsy midbrain tissue from PD patients and age-matched controls were obtained from the ICM Brain Bank and the Neuroceb Pitié-Salpêtrière hostital, Paris. Control brains were obtained from individuals without neurologic or psychiatric disorders. Patients with PD displayed the characteristics of the disease including akinesia, rigidity and/or resting tremor. All patients with PD were treated with dopaminergic agonists and/or L-DOPA and displayed or had displayed dyskinesia. The clinical diagnosis was confirmed by neuropathological examination including the presence of Lewy bodies at least in the substantia nigra. Mean postmortem delay and age at death did not differ between control subjects and patients with PD. A table with details on patients has been appended in supplementary (Supplementary Table S9). SNpc was scraped off the surface of the slides, kept on dry ice under sterile conditions using anatomical landmarks.

### RNA isolation

All the reagents used for RNA isolation and cDNA synthesis were prepared in RNAase free conditions using DEPC water. RNA from mouse brain tissue was isolated using TRIzol reagent (Invitrogen Cat# 15596018) and bromo-chlorophenol (BCP) (Molecular Research Centre, Inc. Cat# BP151)^74^. RNA was isolated from human autopsy brain tissue using the RNeasy Plus Universal Mini Kit (Qiagen Cat# 73404).

### RNA quality check, cDNA library preparation and RNA sequencing

Integrity and purity of RNA samples was determined using Nanodrop, Qubit and BioAnalyzer and samples with Rin value less than 5 were not used. The NEBNext^®^ Ultra™ Directional RNA Library Prep Kit for Illumina^®^ (NEB #E7420G, 30061409) was used for preparing the strand specific libraries from mouse RNA. The sequencing was performed in paired end manner, generating 2X 75 bp long reads and a total of 44-50 million reads._RNA-seq library preparation for human tissue samples was performed using the TruSeq RNA Library Prep Kit v2 (Catalog IDs: RS-122-2001, RS-122-2002). These libraries were prepared in an unstranded manner. The sequencing was performed in paired end manner, generating 2X 100 bp long reads and a total of about 40 million reads. While RNA sequencing on mouse SNpc libraries was performed on Illumina HiSeq2000, RNA-seq on human SNpc libraries was performed on Illumina HiSeq2500.

### Differential gene expression and pathway analysis

Analysis of RNA-seq data was performed on Linux operating system Debian GNU/Linux 8 (Jessie). The raw sequencing data, in the form of FastQ files was checked for quality using FastQC, following which the low quality reads and adapter sequences were removed using Trimmomatic (v0.36)^75^. Further, TopHat2 (v2.1.1)^76^ was used for alignment of reads to mouse or human genomes. Ensembl genome GRCm38 was used as mouse genome reference genome and GRCh37 was used as human reference genome. The resultant alignment files are in the bam (binary alignment/map) format, which were further sorted in the order of names of genes using Samtools, 0.1.19^77^ and finally the numbers of counts for each read in each sample were generated using htseq-count^78^. DESeq2 version 1.22.2^79^ was used for normalization of read counts. DESeq2 analysis was performed on R version 3.5.1.

Normalization by Cufflinks methods^80^ was repeated on the human data after alignment by following the Cufflinks (v2.2.1): Cuffmerge-Cuffdiff pipeline and differentially expressed genes were identified for validation of DESeq2 data.

Data on differential gene expression of developmental genes was also validated by the changing the DESeq2 linear model to account for age, post-mortem interval (PMI) and RIN. The altered model was: ~age+PMI+RIN+condition.

All subsequent data analysis was performed on genes that were differentially expressed and had adjusted p values less than 0.05 as determined using DESeq2 normalization. Functional annotation clusters from differentially expressed genes that were significantly different were created using the Database for Annotation, Visualization and Integrated Discovery (DAVID) bioinformatics resources 6.8^81,82^ and GOplot 1.0.2^83^ was used for visualization of the data. pheatmap 1.0.12^84^ was used for generation of heatmaps and volcano plots were created using EnhancedVolcano 1.5.0^85^. Proteinprotein associations that may result from co-differential expression of genes were drawn using String database^28^. The Venn diagrams were plotted using the tool available at the url: http://bioinformatics.psb.ugent.be/webtools/Venn/.

### Quantitative real time PCR

Human total RNA (200ng) was used for first strand cDNA synthesis using random hexamers, dNTPs and reverse transcriptase from the high capacity cDNA reverse transcription kit (Applied Biosystems Cat# 4368814).

Quantitative real time PCR (qRT-PCR) was performed using SYBR green chemistry for GAP43, RET, LMX1B, EN1 and DAT. The nucleotide sequences for primers used for human tissue gene expression analysis and the qRT-PCR conditions are provided in Supplementary Tables S10 and S11. Two endogenous controls, namely β-actin and GAPDH were used for normalization. For mouse SNpc samples, qRT-PCR was performed using SYBR green chemistry for Dat, Th, Lmx1b, En1, Ret and Gap43. The nucleotide sequences for primers used for mouse tissue gene expression analysis and the qRT-PCR conditions are provided in Supplementary Table S12. 18S rRNA was used for normalization.

### LMX1B over expression construct

Expression construct for human LMX1B was subcloned from the tetO-ALN plasmid^86^. tetO-ALN was a gift from John Gearhart (Addgene plasmid # 43918; http://n2t.net/addgene:43918; RRID: Addgene_43918). The open reading frame (ORF) for human LMX1B was PCR-amplified using the following primer sequences: Forward primer: TAAGCATCTAGAATGTTGGACGGCATCAAG; Reverse primer: TGCTTAGGATCCTCAGGAGGCGAAGTAGGAACT. The ORF thus PCR-amplified was cloned into the pcDNA™3.1/V5-His TOPO™ TA vector (ThermoFisher Scientific, Cat # K480040) following manufacturer’s protocol.

### Experiments with SH-SY5Y cells

SH-SY5Y cells (ATCC^®^ CRL-2266™) were cultured in Minimum Essential Medium Eagle (MEM; Sigma, Cat# M0643) that was supplemented with 10% v/v fetal bovine serum (FBS; Gibco, Cat# 26140079, Origin: U.S) and 1X penicillin-streptomycin (Gibco, Cat# 15070063). SH-SY5Y cells were plated at a confluency of 70% on glass cover slips (VWR, Cat# 26022) that were acid washed and precoated with poly-D-lysine (0.1 mg/ml) in 12-well plates.

Within ~16 hours of plating, SH-SY5Y cells were transfected with pcDNA3.1-LMX1B using Lipofectamine™ LTX Reagent with PLUS™ Reagent (Invitrogen, Cat # 15338030) in OptiMEM medium (Gibco, Cat# 22600-050). The media was changed to MEM with FBS and penicillin-streptavidin after 6 hours.

SH-SY5Y cells were treated with MPP^+^ (1mM; Sigma-Aldrich D048) 48 hours after transfection with pcDNA3.1-LMX1B. The cells were washed with phosphate buffered saline (PBS) and fixed with 2% paraformaldehyde (w/v) 24 hours after MPP^+^ exposure. For quantification of LMX1B overexpression, RNA was isolated from the transfected cells using the RNeasy Plus Micro kit (Qiagen Cat# 74034).

The extent of cell death following MPP^+^ treatment with and without LMX1B overexpression was assessed by using the terminal deoxynucleotidyl transferase mediated dUTP-biotin nick end-labeling (TUNEL) method (In situ cell death detection kit, TMR red, Roche, Cat# 12156792910), according to the manufacturer’s protocol. Ten fields were imaged from three independent coverslips for each experimental condition. Analysis was performed to quantify the average number of TUNEL-positive (Red) cells and was blinded.

### Statistics

Samples were analyzed in triplicates. Relative gene expression was analyzed using ΔΔCt method. Data was analyzed and represented using Graphpad Prism (Graphpad Prism Inc, USA). Statistical significance was determined using Student’s *t* test. All data are presented as mean ± SEM.

### Study Approval

Animal experiments were approved by the institutional animal ethical review board, named ‘Institutional Animal and Ethics Committee’ of Indian Institute of Science (Protocol# CAF/Ethics/267/2012). Animals were handled according to the guidelines of Committee for the Purpose of Control and Supervision of Experiments on Animals (CPCSEA), Government of India.

All experiments on human autopsy brain tissue were carried out following approval from Institutional Human Ethics Committee of the Indian Institute of Science and all guidelines were followed (IHEC approval# 10/1/2015).

## Acknowledgements

The study was funded by TATA Trusts (VR). AV received research fellowship from Council of Scientific and Industrial Research (CSIR), Government of India. EH acknowledges the financial support from CNRS, INSERM, ICM, UPMC and from the program “Investissements d’avenir” ANR-10-IAIHU-06. The authors declare no other financial disclosures. The authors are grateful to Pr Charles Duyckaerts for the neuropathological examination Annick Prigent from the ICM Histomics platform and Sabrina Leclere-Turbant from Neuroceb for their help in preparing the human postmortem samples.

## Authors’ Contributions

Research project was conceptualized by VR, organized by AV, EH and VR and executed by AV, PS, and BG. Statistical analysis was designed and executed by AV, and reviewed and critiqued upon by EH and VR. First draft of the manuscript was written by AV and VR and was further reviewed and critiqued upon by EH.

## References

1. Parkinson, J. D. An Essay on the Shaking Palsy. London Whittingham Rowl. Sherwood, Neely, Jones 14, 223–236 (1817).

2. Cotzias, G. C. L-Dopa for Parkinsonism. N. Engl. J. Med. 278, 630 (1968).

3. Hornykiewicz, O. A brief history of levodopa. J. Neurol. 257, S249–52 (2010).

4. Ascherio, A. & Schwarzschild, M. A. The epidemiology of Parkinson’s disease: risk factors and prevention. Lancet Neurol. 15, 1257–1272 (2016).

5. Schapira, A. H. V. et al. Mitochondrial Complex I Deficiency in Parkinson’s Disease. J. Neurochem. 54, 823–827 (1990).

6. Park, J., Kim, Y. & Chung, J. Mitochondrial dysfunction and Parkinson’s disease genes: insights from Drosophila. Dis. Model. Mech. 2, (2009).

7. Sherman, M. Y. & Goldberg, A. L. Cellular Defenses against Unfolded Proteins: A Cell Biologist Thinks about Neurodegenerative Diseases. Neuron 29, 15–32 (2001).

8. Carvalho, A. N. et al. Ubiquitin–Proteasome System Impairment and MPTP-Induced Oxidative Stress in the Brain of C57BL/6 Wild-type and GSTP Knockout Mice. Mol. Neurobiol. 47, 662–672 (2013).

9. Stefanis, L. α-Synuclein in Parkinson’s disease. Cold Spring Harb. Perspect. Med. 2, a009399–a009399 (2012).

10. Hirsch, E. C., Vyas, S. & Hunot, S. Neuroinflammation in Parkinson’s disease. Parkinsonism Relat. Disord. 18, S210–S212 (2012).

11. Pifl, C. et al. Is Parkinson’s Disease a Vesicular Dopamine Storage Disorder? Evidence from a Study in Isolated Synaptic Vesicles of Human and Nonhuman Primate Striatum. J. Neurosci. 34, (2014).

12. Sofic, E. et al. Increased iron (III) and total iron content in post mortem substantia nigra of parkinsonian brain. J. Neural Transm. 74, 199–205 (1988).

13. Ayton, S. & Lei, P. Nigral iron elevation is an invariable feature of Parkinson’s disease and is a sufficient cause of neurodegeneration. Biomed Res. Int. 2014, 581256 (2014).

14. Borrageiro, G., Haylett, W., Seedat, S., Kuivaniemi, H. & Bardien, S. A review of genome-wide transcriptomics studies in Parkinson’s disease. Eur. J. Neurosci. 47, 1–16 (2018).

15. Sutherland, G. T. et al. A Cross-Study Transcriptional Analysis of Parkinson’s Disease. PLoS One 4, e4955 (2009).

16. Greene, J. G. Current status and future directions of gene expression profiling in Parkinson’s disease. Neurobiol. Dis. 45, 76–82 (2012).

17. Simunovic, F. et al. Gene expression profiling of substantia nigra dopamine neurons: further insights into Parkinson’s disease pathology. Brain 132, 1795–1809 (2009).

18. Miller, R. M. & Federoff, H. J. Altered Gene Expression Profiles Reveal Similarities and Differences Between Parkinson Disease and Model Systems. Neurosci. 11, 539–549 (2005).

19. Zhang, Y., James, M., Middleton, F. A. & Davis, R. L. Transcriptional analysis of multiple brain regions in Parkinson’s disease supports the involvement of specific protein processing, energy metabolism, and signaling pathways, and suggests novel disease mechanisms. Am. J. Med. Genet. Part B Neuropsychiatr. Genet. 137B, 5–16 (2005).

20. Malone, J. H. & Oliver, B. Microarrays, deep sequencing and the true measure of the transcriptome. BMC Biol. 9, 34 (2011).

21. Kogenaru, S., Qing, Y., Guo, Y. & Wang, N. RNA-seq and microarray complement each other in transcriptome profiling. BMC Genomics 13, 629 (2012).

22. Saeed, U. et al. Redox Activated MAP Kinase Death Signaling Cascade Initiated by ASK1 is not Activated in Female Mice Following MPTP: Novel Mechanism of Neuroprotection. Neurotox. Res. 16, 116–126 (2009).

23. Maraganore, D. M. et al. UCHL1 is a Parkinson’s disease susceptibility gene. Ann. Neurol. 55, 512–521 (2004).

24. Cook, C. & Petrucelli, L. A critical evaluation of the ubiquitin-proteasome system in Parkinson’s disease. Biochim. Biophys. Acta 1792, 664–75 (2009).

25. Grima, B. et al. A single human gene encoding multiple tyrosine hydroxylases with different predicted functional characteristics. Nature 326, 707–711 (1987).

26. Lewis, D. A., Melchitzky, D. S. & Haycock, J. W. Four isoforms of tyrosine hydroxylase are expressed in human brain. Neuroscience 54, 477–492 (1993).

27. Iwata, N., Kobayashi, K., Sasaoka, T., Hidaka, H. & Nagatsu, T. Structure of the mouse tyrosine hydroxylase gene. Biochem. Biophys. Res. Commun. 182, 348–354 (1992).

28. Szklarczyk, D. et al. STRING v11: protein-protein association networks with increased coverage, supporting functional discovery in genome-wide experimental datasets. Nucleic Acids Res. 47, D607–D613 (2019).

29. Riley, B. E. et al. Systems-Based Analyses of Brain Regions Functionally Impacted in Parkinson’s Disease Reveals Underlying Causal Mechanisms. PLoS One 9, e102909 (2014).

30. Keo, A. et al. Transcriptomic signatures of brain regional vulnerability to Parkinson’s disease. Commun. Biol. 3, 101 (2020).

31. Blaudin de Thé, F.-X., Rekaik, H., Prochiantz, A., Fuchs, J. & Joshi, R. L. Neuroprotective Transcription Factors in Animal Models of Parkinson Disease. Neural Plast. 2016, 1–11 (2016).

32. Domanskyi, A., Alter, H., Vogt, M. A., Gass, P. & Vinnikov, I. A. Transcription factors Foxa1 and Foxa2 are required for adult dopamine neurons maintenance. Front. Cell. Neurosci. 8, 275 (2014).

33. Alavian, K. N. et al. The lifelong maintenance of mesencephalic dopaminergic neurons by Nurr1 and engrailed. J. Biomed. Sci. 21, 27 (2014).

34. Lei, Z., Jiang, Y., Li, T., Zhu, J. & Zeng, S. Signaling of Glial Cell Line-Derived Neurotrophic Factor and Its Receptor GFRα1 Induce Nurr1 and Pitx3 to Promote Survival of Grafted Midbrain-Derived Neural Stem Cells in a Rat Model of Parkinson Disease. J. Neuropathol. Exp. Neurol. 70, 736–747 (2011).

35. Jacobs, F. M. J. et al. Identification of Dlk1, Ptpru and Klhl1 as novel Nurr1 target genes in meso-diencephalic dopamine neurons. Development 136, 2363–2373 (2009).

36. Thuret, S., Bhatt, L., O’Leary, D. D.. & Simon, H. H. Identification and developmental analysis of genes expressed by dopaminergic neurons of the substantia nigra pars compacta. Mol. Cell. Neurosci. 25, 394–405 (2004).

37. Kramer, E. R. & Liss, B. GDNF-Ret signaling in midbrain dopaminergic neurons and its implication for Parkinson disease. FEBS Lett. 589, 3760–3772 (2015).

38. Drinkut, A. et al. Ret is essential to mediate GDNF’s neuroprotective and neuroregenerative effect in a Parkinson disease mouse model. Cell Death Dis. 7, e2359 (2016).

39. Murakami, M. et al. Sphingosine kinase 1/S1P pathway involvement in the GDNF-induced GAP43 transcription. J. Cell. Biochem. 112, 3449–3458 (2011).

40. Murakami, M. et al. RET signaling-induced SPHK1 gene expression plays a role in both GDNF-induced differentiation and MEN2-type oncogenesis. J. Neurochem. 102, 1585–1594 (2007).

41. Simon, H. H., Bhatt, L., Gherbassi, D., Sgado, P. & Alberi, L. Midbrain dopaminergic neurons: determination of their developmental fate by transcription factors. Ann. N. Y. Acad. Sci. 991, 36–47 (2003).

42. Prakash, N. & Wurst, W. Genetic networks controlling the development of midbrain dopaminergic neurons. J. Physiol. 575, 403–410 (2006).

43. Abeliovich, A. & Hammond, R. Midbrain dopamine neuron differentiation: Factors and fates. Dev. Biol. 304, 447–454 (2007).

44. Bissonette, G. B. & Roesch, M. R. Development and function of the midbrain dopamine system: what we know and what we need to. 15, 62–73 (2017).

45. Hegarty, S. V., Sullivan, A. M. & O’Keeffe, G. W. Midbrain dopaminergic neurons: A review of the molecular circuitry that regulates their development. Dev. Biol. 379, 123–138 (2013).

46. Arenas, E., Denham, M. & Villaescusa, J. C. How to make a midbrain dopaminergic neuron. Development 142, 1918–36 (2015).

47. Smidt, M. P. Molecular Programming of Mesodiencephalic Dopaminergic Neuronal Subsets. Front. Neuroanat. 11, 59 (2017).

48. Hoekstra, E. J. et al. Lmx1a Encodes a Rostral Set of Mesodiencephalic Dopaminergic Neurons Marked by the Wnt/B-Catenin Signaling Activator R-spondin 2. PLoS One 8, e74049 (2013).

49. Hoekstra, E. J. et al. *Lmx1a* is an activator of *Rgs4* and *Grb10* and is responsible for the correct specification of rostral and medial mdDA neurons. Eur. J. Neurosci. 37, 23–32 (2013).

50. Baek, S., Choi, H. & Kim, J. Ebf3-miR218 regulation is involved in the development of dopaminergic neurons. Brain Res. 1587, 23–32 (2014).

51. Kouwenhoven, W. M., Von Oerthel, L. & Smidt, M. P. Pitx3 and En1 determine the size and molecular programming of the dopaminergic neuronal pool. PLoS One 12, 1–18 (2017).

52. Veenvliet, J. V. et al. Specification of dopaminergic subsets involves interplay of En1 and Pitx3. Development 140, 4116–4116 (2013).

53. Villaescusa, J. C. et al. A PBX1 transcriptional network controls dopaminergic neuron development and is impaired in Parkinson’s disease. EMBO J. 35, 1963–78 (2016).

54. Peng, C. et al. Pitx3 Is a Critical Mediator of GDNF-Induced BDNF Expression in Nigrostriatal Dopaminergic Neurons. J. Neurosci. 31, 12802–12815 (2011).

55. Alavian, K. N. et al. The lifelong maintenance of mesencephalic dopaminergic neurons by Nurr1 and engrailed. J. Biomed. Sci. 21, 27 (2014).

56. Lee, H. S. et al. Foxa2 and Nurr1 synergistically yield A9 nigral dopamine neurons exhibiting improved differentiation, function, and cell survival. Stem Cells 28, 501–512 (2010).

57. Lin, W. et al. Foxa1 and Foxa2 function both upstream of and cooperatively with Lmx1a and Lmx1b in a feedforward loop promoting mesodiencephalic dopaminergic neuron development. Dev. Biol. 333, 386–396 (2009).

58. Simon, H. H., Saueressig, H., Wurst, W., Goulding, M. D. & OLeary, D. D. M. Fate of Midbrain Dopaminergic Neurons Controlled by the Engrailed Genes. J. Neurosci. 21, 3126 LP–3134 (2001).

59. Laguna, A. et al. Dopaminergic control of autophagic-lysosomal function implicates Lmx1b in Parkinson’s disease. Nat Neurosci 18, 826–835 (2015).

60. Doucet-Beaupré, H. et al. Lmx1a and Lmx1b regulate mitochondrial functions and survival of adult midbrain dopaminergic neurons. Proc. Natl. Acad. Sci. 113, E4387–E4396 (2016).

61. Greenwood, T. A. et al. Genome-wide Association of Endophenotypes for Schizophrenia From the Consortium on the Genetics of Schizophrenia (COGS) Study. JAMA psychiatry (2019). doi:10.1001/jamapsychiatry.2019.2850

62. Consortium, S. W. G. of the P. G. Biological insights from 108 schizophrenia-associated genetic loci. Nature 511, 421–427 (2014).

63. Evangelou, E. et al. New alcohol-related genes suggest shared genetic mechanisms with neuropsychiatric disorders. Nat. Hum. Behav. 3, 950–961 (2019).

64. Lane, J. M. et al. Genome-wide association analyses of sleep disturbance traits identify new loci and highlight shared genetics with neuropsychiatric and metabolic traits. Nat. Genet. 49, 274–281 (2017).

65. Zhou, H. et al. Genetic Risk Variants Associated With Comorbid Alcohol Dependence and Major Depression. JAMA psychiatry 74, 1234–1241 (2017).

66. Brazel, D. M. et al. Exome Chip Meta-analysis Fine Maps Causal Variants and Elucidates the Genetic Architecture of Rare Coding Variants in Smoking and Alcohol Use. Biol. Psychiatry 85, 946–955 (2019).

67. Jansen, P. R. et al. Genome-wide analysis of insomnia in 1,331,010 individuals identifies new risk loci and functional pathways. Nat. Genet. 51, 394–403 (2019).

68. Spada, J. et al. Genome-wide association analysis of actigraphic sleep phenotypes in the LIFE Adult Study. J. Sleep Res. 25, 690–701 (2016).

69. Sherva, R. et al. Genome-wide Association Study of Cannabis Dependence Severity, Novel Risk Variants, and Shared Genetic Risks. JAMA psychiatry 73, 472–480 (2016).

70. Matoba, N. et al. GWAS of smoking behaviour in 165,436 Japanese people reveals seven new loci and shared genetic architecture. Nat. Hum. Behav. 3, 471–477 (2019).

71. Jackson-Lewis, V. & Przedborski, S. Protocol for the MPTP mouse model of Parkinson’s disease. Nat. Protoc. 2, 141–151 (2007).

72. Gundersen, H. J. & Jensen, E. B. The efficiency of systematic sampling in stereology and its prediction. J. Microsc. 147, 229–63 (1987).

73. West, M. J., Slomianka, L. & Gundersen, H. J. G. Unbiased stereological estimation of the total number of neurons in the subdivisions of the rat hippocampus using the optical fractionator. Anat. Rec. 231, 482–497 (1991).

74. Chomczynski, P. & Sacchi, N. The single-step method of RNA isolation by acid guanidinium thiocyanate–phenol–chloroform extraction: twenty-something years on. Nat. Protoc. 1, 581–585 (2006).

75. Bolger, A. M., Lohse, M. & Usadel, B. Trimmomatic: a flexible trimmer for Illumina sequence data. Bioinformatics 30, 2114–20 (2014).

76. Kim, D. et al. TopHat2: accurate alignment of transcriptomes in the presence of insertions, deletions and gene fusions. Genome Biol. 14, R36 (2013).

77. Li, H. et al. The Sequence Alignment/Map format and SAMtools. Bioinformatics 25, 2078–2079 (2009).

78. Anders, S., Pyl, P. T. & Huber, W. HTSeq--a Python framework to work with high-throughput sequencing data. Bioinformatics 31, 166–9 (2015).

79. Love, M. I., Huber, W. & Anders, S. Moderated estimation of fold change and dispersion for RNA-seq data with DESeq2. Genome Biol. 15, 550 (2014).

80. Trapnell, C. et al. Differential gene and transcript expression analysis of RNA-seq experiments with TopHat and Cufflinks. Nat. Protoc. 7, 562–78 (2012).

81. Huang, D. W., Sherman, B. T. & Lempicki, R. A. Bioinformatics enrichment tools: paths toward the comprehensive functional analysis of large gene lists. Nucleic Acids Res. 37, 1–13 (2009).

82. Huang, D. W., Sherman, B. T. & Lempicki, R. A. Systematic and integrative analysis of large gene lists using DAVID bioinformatics resources. Nat. Protoc. 4, 44–57 (2009).

83. Walter, W., Sanchez-Cabo, F. & Ricote, M. GOplot: an R package for visually combining expression data with functional analysis. Bioinformatics 31, 2912–2914 (2015).

84. Kolde, R. Pheatmap: pretty heatmaps. R Packag. version 61, 617 (2012).

85. Blighe, K., Rana, S. & Lewis, M. EnhancedVolcano: Publication-ready volcano plots with enhanced colouring and labeling. (2019).

86. Addis, R. C. et al. Efficient conversion of astrocytes to functional midbrain dopaminergic neurons using a single polycistronic vector. PLoS One 6, e28719–e28719 (2011).

