## SupplementaryFig1 for "Genes critical for development and differentiation of dopaminergic neurons are downregulated in Parkinson’s disease" (from another version of this preprint: version 2, 2020-04-09)

Address for correspondence

**Supplementary data**

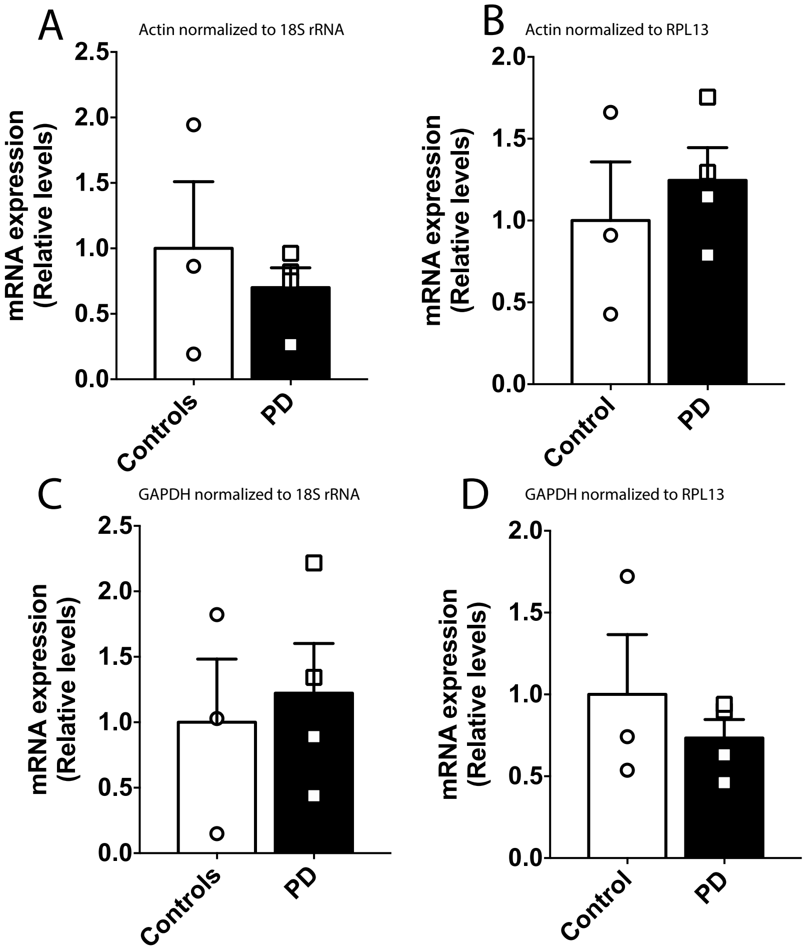


**Figure 1: Lack of differential gene expression of β-actin and GAPDH in human PD.** mRNA expression of β-actin does not change in PD human autopsy tissue as investigated through qRT-PCR when normalized to either 18S rRNA (A) or to RPL13 (B). The expression of GAPDH also does not alter in PD upon normalization to 18S rRNA (C) and RPL13 (D). Each point represents an individual sample. For controls n=3, for human PD samples, n=4. Unpaired, two-tailed Student' t test has been performed on each control versus PD comparison. Data has been presented as mean ± SEM.
